# Impact of exercise intervention on IGF-1 signaling related to muscle regeneration and physical performance in aged mice

**DOI:** 10.1101/2025.03.14.643197

**Authors:** Taewan Kim, Jinkyung Cho, Yoonhwan Kim, Joohyung Kim, Sanggu Woo, Donghyun Kim

**Author notes:** Corresponding authors’ information: Donghyun Kim, Ph.D., Department of Sports and Health Science, Hanbat National University, 125, Dongseo-daero, Yuseong-gu, Daejeon, Republic of Korea.

## Abstract

Aging encompasses the natural processes of birth, growth, and aging, during which the functional ability of muscles gradually decreases, leading to the loss of muscle size and reduced exercise performance known as sarcopenia. This condition is closely associated with weakness, osteoporosis, and degenerative diseases, increasing the risk of falls, fractures, metabolic diseases, and mortality due to limitations in physical performance among the elderly. This study investigated the effects of exercise intervention on biological markers related to skeletal muscle mass and functions in conjunction with aging. At age of four or twenty, the C57BL/6 mice were assigned to Young control (Y-Con, n = 10) or exercise training (Y--Exe, n = 10), and Aged control (A-Con, n=10) or exercise training (A-Exe, n = 10). Exercise intervention was performed on a rodent motor-driven treadmill with a frequency of 5 days per week for 8 weeks. As a consequence, exercise intervention in mice resulted in positive changes in IGF-1 signaling and muscle phenotype compared to mice that did not undergo exercise intervention, specifically showing prominent effects in the A-Exe group compared to the A-Con group. The mitigating effects of exercise intervention on age-related skeletal muscle dysfunction were accompanied by enhanced exercise performance and muscle function, as assessed by grip strength and the rotarod test. The current findings support previous studies that have reported the positive effect of exercise intervention in alleviating age-related declines in exercise performance and muscle function in older adults.

## Introduction

Aging is associated with a progressive decline in skeletal muscle mass and regenerative capacity. Impaired regeneration process results in weakened muscles strength, which will limit the physical function and affect life quality ^1^. Sarcopenia occurs commonly as an age-related process in older people, progressive loss of skeletal muscle mass and function, which is accompanied by reduced motor nerve cells ^2, 3^ and satellite cells ^4^, mitochondrial dysfunction ^5^, elevation of oxidative stress ^6^. Some studies indicate that age-related physiological changes such as sarcopenia leads to functional limitations, impaired mobility, disability, falls, and fractures, which in turn result in the loss of independence, frailty, and an increased risk of mortality ^7, 8^. Despite the rapid increase in the elderly population, the molecular mechanisms of sarcopenia remain incompletely understood, and targeted medications for preventing this condition are unclear and ineffective.

Many epidemiological studies have reported that the physiological changes induced by exercise training effectively control the incidence of sarcopenia by enhancing muscle strength ^9^, improving body composition ^10^, and inducing cardiovascular ^11^ and nervous system adaptations ^12^. Exercise-triggered peripheral adaptations may be related to multiple intracellular signaling systems that arise during the process of muscle regeneration and growth, which can improve aged-related muscle atrophy ^13, 14^. Particularly, the IGF-1/AKT/mTOR signaling system has been closely associated with regulating age-related changes in skeletal muscles ^15, 16^. The findings of clinical and experimental studies demonstrate that the IGF-1 signaling pathway is independently associated with the reduction of skeletal muscle mass ^17^, inhibition of age-related muscle wasting and weakness ^18^, and modulating mitochondrial function, ROS detoxification, and the basal inflammatory state occurring at old age ^19^. In addition, IGF-1 also potentiates skeletal muscle regeneration via the activation of skeletal muscle stem (satellite) cells, which may contribute to muscle hypertrophy and/or inhibit atrophy ^20^. For this reason, exercise-induced metabolic changes contribute to the regeneration and recovery of damaged skeletal muscles, thereby preventing or slowing the progression of sarcopenia and related functional decline in the elderly ^21^.

Therefore, we conducted a study to investigate the expression levels of IGF-1 signaling biomarkers along with the myogenesis-related proteins in the gastrocnemius. This study aimed to explore the role of the IGF-1 signaling pathway in modulating the physiological responses of muscle to exercise and aging in naturally aging mice.

## Materials and methods

### Animal

We used 4-month-old and 20-month-old C57/BL6N mice purchased from Orient Bio. These mice were classified into young (weight:20.7 ± 1.8 g) and old (weight:34.2 ±2.9 g) groups. All mice were housed in sterile cages under a day-and-night cycle and had free access to solid food (sterilized using high-pressure steam) and drinking water during the experimental period. The temperature was maintained at 22–24 °C and the humidity was 50 ± 10%.

### Experimental Protocol

The experimental animals were randomly divided into four groups:1) Young control group (Y-Con, n = 10), 2) Young exercise group (Y-Exe, n = 10), 3) Aged control group (A-Con, n = 10), and 4) Aged exercise group (A-Exe, n = 10). All animals were allowed to eat ad libitum and body weight was recorded twice a week for the duration of the study. The Exe groups were trained on a motor-driven treadmill (Columbus Instruments, Inc., Columbus, OH, USA) with duration of 40 min per session and a frequency of 5 days per week for 8 weeks. The mice ran on the treadmill at zero inclination and a speed of 15 m/min for 1-2 weeks and 20 m/min for 3-8 weeks. After the 8 week exercise treatment, grip strength and rotarod tests were performed to assess exercise performance.

### Grip strength test

All mice underwent weekly muscle strength measurements using a muscle strength tester (BIOSSEB’s Grip strength test, France), commencing from the first week of the experiment. The measurement procedure included an experimenter carefully assisting the mouse by holding its torso and tail, directing it to grasp the bar of the muscle strength meter located 10 cm from the ground. Subsequently, the tail was steadily pulled back, causing the paws to release from the bar. The peak muscle strength (g) was recorded, and the procedure was repeated twice to obtain an average tension.

### Rotarod test

The motor performance of all mice was assessed using a Rota-Rod test apparatus (mouse Rota-Rod 47700, UGO BASILE, Milano, Italy). The procedure involved placing the mouse on a rotating cylindrical bar, initially rotating at a rate of 4 rpm and gradually accelerating. The rotation speed continuously increased, reaching a maximum of 40 rpm within 5 minutes. The time (seconds) until the mouse fell was then measured. Measurements were conducted for up to 300 seconds, with readings taken once every 2 weeks for 8weeks. Average values were calculated from two consecutive trials separated by a 20 minute interval.

### Hematoxylin & Eosin Staining

The muscle tissues extracted from the gastrocnemius were placed in 4% neutral buffered formalin (NBF) solution and fixed at 4 °C for 24 h. The fixed muscle tissues were then processed in 70%, 80%, 90%, and 100% ethanol solutions for more than 16 h, and then fixed in a paraffin block. Using a paraffin section (Leica, Nussloch, Germany), sections of 4 μm thickness were placed on glass slides. The glass slides were dehydrated by first treating them in xylene solution for 3 min and then soaking in 100%, 90%, 70%, and 50% ethanol for 5 min each. The slides were then washed with distilled water (DW), stained with hematoxylin for 5 min, washed several times with DW, stained with eosin for 2 min, washed several times with DW, and soaked in 50%, 70%, 90%, and 100% ethanol for 5 min to dehydrate them. The slides were then immersed in xylene thrice for 3 min each and covered with a cover slide and mounting solution. The images were captured by a Leica DM759 light micorosope and analysed using Image J software.

### Western blot

Western blot analysis was performed to evaluate protein expression in the gastrocnemius muscle. The extracted gastrocnemius tissues were lysed in lysis buffer containing 50 mM HEPES (pH 7.5), 150 mM NaCl, 10% glycerol, 1% Triton X-100, 1.5 mM magnesium chloride hexahydrate, 1 mM phenylmethylsulfonyl fluoride (PMSF), 2 mg/mL leupeptin, 1 mg/mL pepstatin, 1 mM sodium orthovanadate, and 100 mM sodium fluoride. The supernatant was collected by centrifugation at 14,000 rpm for 15 min, and the protein concentration was quantified using the Bradford assay. Total protein concentrations of 20 μg for each group were electrophoresed on a 10% SDS-polyacrylamide gel, transferred to a nitrocellulose membrane, and incubated at room temperature for 1 h with 4% skim milk, followed by incubation with the primary antibody for 24 h at 4 °C. The primary antibodies used were rabbit anti-IGF1 (1:1000; ab9572; Abcam, Cambridge), rabbit anti-p-PI3K (1:1000; #4228; Cell Signaling, Danvers, MA), rabbit anti-PI3K (1:1000; ab302958; Abcam, Cambridge), rabbit anti-p-AKT (1:1000; 44-621G; Thermo Fisher Scientific, Waltham, MA), rabbit anti-AKT (1:1000; #4060; Cell Signaling), rabbit anti-β-actin (1:1000; A300-491A; Bethyl, TX, USA), rabbit anti-p-mTOR (1:1000; #5536; Cell Signaling), rabbit anti-mTOR (1:1000; ab134903; Abcam), rabbit anti-p-p70S6K (1:1000; #97596; Cell Signaling), rabbit anti-p70S6K (1:1000; # 34475; Cell Signaling). Subsequently, the membrane was washed thrice for 10 min each with 0.05% Tween 20 in 1X TBST and then incubated with horseradish peroxidase (HRP)-conjugated secondary antibodies for 30 min at room temperature. After 3 washes with 0.05% Tween 20 in 1X TBST for 10 min, the membrane was developed using chemiluminescent (ECL) western blotting detection reagents (Amersham, Pharmacia Biotech GmbH, Freiburg, Germany). Protein expression was measured using an Image-Pro Plus image analyzer (Media Cybernetics Inc., Silver Spring, MD, USA).

### Statistical analyses

Data are presented as means ± SD. Two-way ANOVA was used to test the main effects of time (Grip strength test: 1 week, 2 weeks, 3 weeks, 4 weeks, 5 weeks, 6 weeks, 7 weeks, 8 weeks, Rotarod test: 2week, 4week, 6week, 8week) and group (Y-Con, Y-Exe, A-Con, A-Exe), as well as the group by time interaction on grip strength in the grip strength test and latency to fall in the rotarod test. One-way ANOVAs were used to compare any significant differences in the outcome variables between the groups of mice. The Tukey post-hoc multiple comparison tests were used to test main effects (two-way ANOVA) and group differences (one way ANOVA) in the measured parameters, if necessary. Statistical significance was set at p<0.05 with the SPSS-PC software (version 23.0).

## Results

The current study was conducted using young and old C57BL/6N wild-type mice to investigate the effects of 8 weeks of endurance exercise on muscle regeneration, the IGF-1 signaling pathway, and exercise performance during aging.

### Exercise intervention showed positive effects on improving muscle strength in aged mice

To explore the changes in muscle strength induced by treadmill exercise, grip strength was measured weekly (Fig 1). There was a significant increase in muscle strength in exercise training mice across both group and time. Furthermore, there was main effect for time (*p* < 0.001) or group (*p* < 0.001), there was a significant group by time interaction (*p* < 0.001). Post hoc analysis revealed that both exercise-trained mice such as Y-Exe and A-Exe showed a significant increase in final grip strength test at 8^th^ week (Y-con, *p* < 0.001; A-Con, *p* < 0.001; A-Exe, *p* < 0.001 vs Y-Exe, and Y-con, *p* < 0.001; A-Con, *p* < 0.001 vs A-Exe).

**Fig 1.**
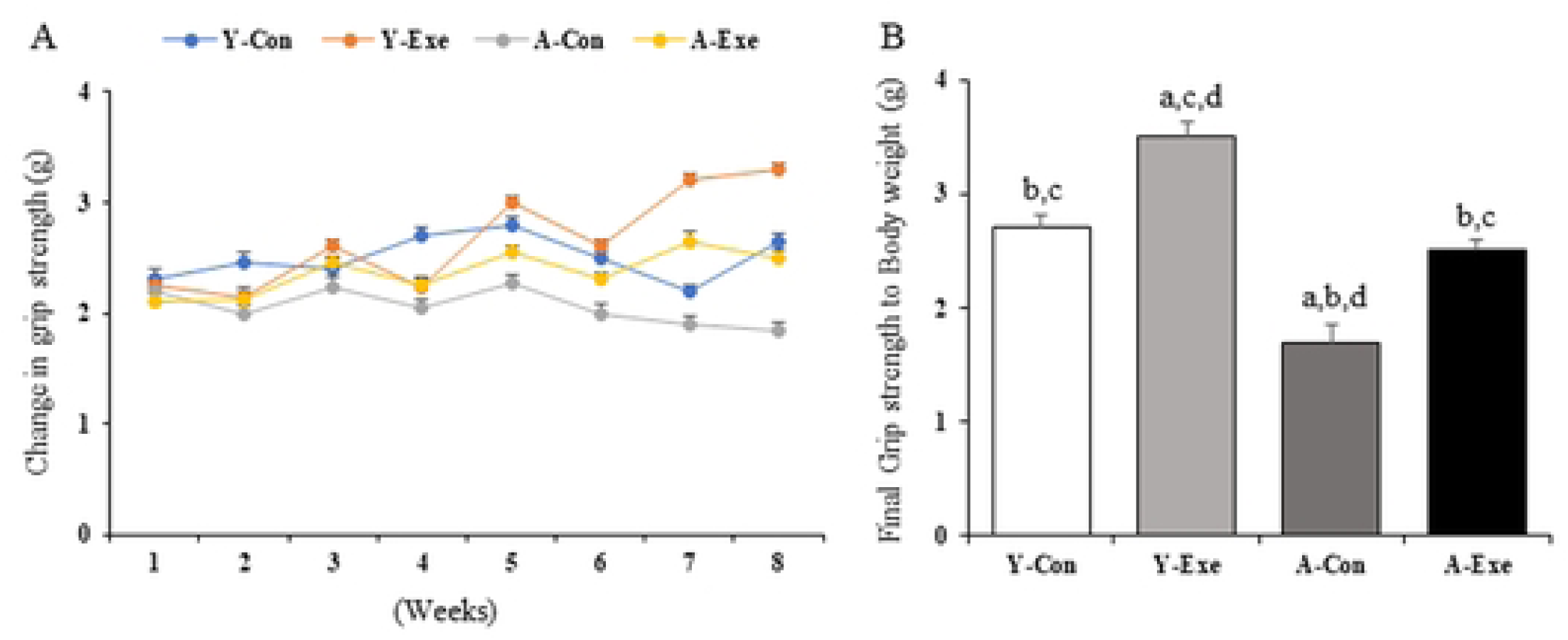
Change in grip strength (A), Final grip strength to Body weight (B) in hindlimb from young or old C57BL/6 mice. Y-Con; Young-Control, Y-Exe; Young-Exercise, A-Con; Aged-Control, A-Exe; Aged-Exercise. (a) *p* < 0.05 vs Y-Con, (b) *p* < 0.05 vs Y-Exe, (c) *p* < 0.05 vs A-Con, (d) *p* < 0.05 vs A-Exe, Values are presented a mean±SD; n= 10 per each group.

### Exercise intervention showed positive effects on improving motor performance in aged mice

The rotarod test was performed to assess locomotion and balance, which were associated with age-related coordination deficits. The test measured the latency to fall, assessing the time taken for the animal to drop off the rotating rod (Fig 2). There was a significant increase in latency to fall in exercise training mice across both groups (*p* = 0.001), and time (*p* = 0.001). Furthermore, there was no main effect for time or group, there was a significant group by time interaction (*p* = 0.001). Post hoc analysis revealed that both exercise-trained mice such as Y-Exe and A-Exe showed a significant increase in final latency to fall at 8th week (Y-con, *p* < 0.001; A-Con, *p* < 0.001; A-Exe, p < 0.001 vs Y-Exe, and Y-con, *p* = 0.45; A-Con, *p* = 0.001 vs A-Exe).

**Fig 2.**
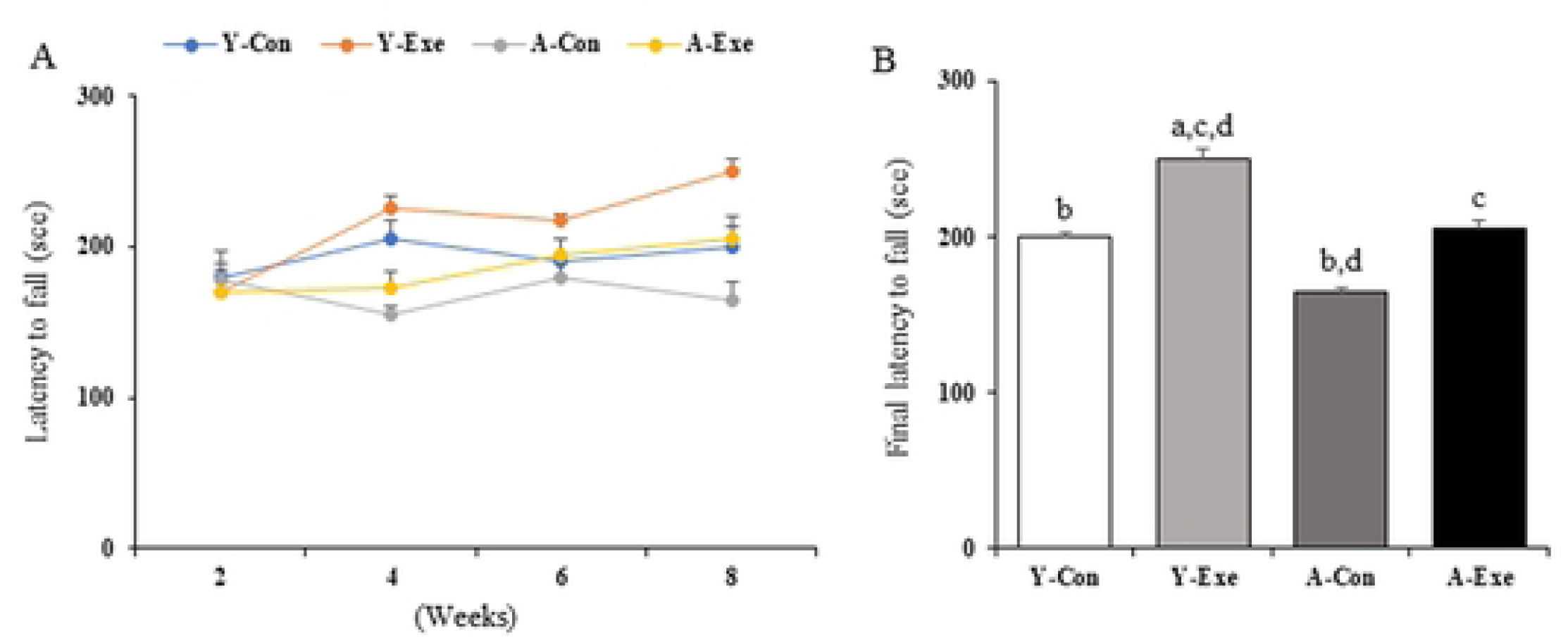
Latency to fall (A), Final Latency to fall (B) from young or old C57BL/6 mice. Y-Con; Young-Control, Y-Exe; Young-Exercise, A-Con; Aged-Control, A-Exe; Aged-Exercise. (a) *p* < 0.05 vs Y-Con, (b) *p* < 0.05 vs Y-Exe, (c) *p* < 0.05 vs A-Con, (d) *p* < 0.05 vs A-Exe, Values are presented a mean±SD; n= 10 per each group

### Exercise intervention did not show positive effects on muscle fiber size in aged mice

Regarding alterations in muscle fiber size, the size of muscle fibers in each group is shown in Fig 3. The changes in muscle fiber size were greater in young mice than in aged mice. Y-Exe mice had largest muscle fiber size, whereas A-Con mice had the smallest muscle fiber size among all groups. Although there was an increase in muscle fiber size observed in both young and aged mice, these differences did not reach statistical significance.

**Fig 3.**
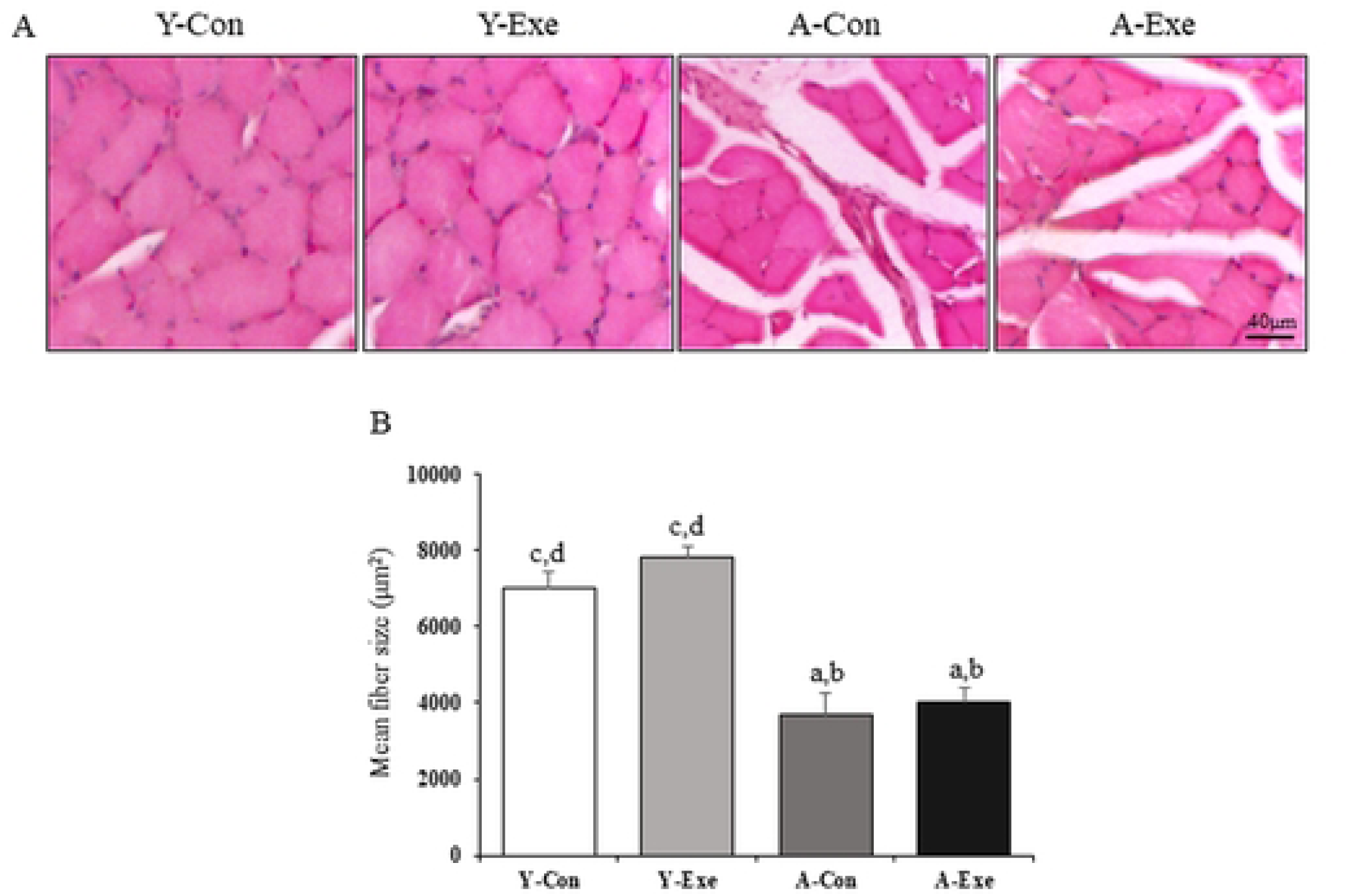
H&E staining (A), Mean fiber size (B) from young or old C57BL/6 mice. Y-Con; Young-Control, Y-Exe; Young-Exercise, A-Con; Aged-Control, A-Exe; Aged-Exercise. (a) *p* < 0.05 vs Y-Con, (b) *p* < 0.05 vs Y-Exe, (c) *p* < 0.05 vs A-Con, (d) *p* < 0.05 vs A-Exe, Values are presented a mean±SD; n= 10 per each group.

### Exercise intervention showed positive effects on improving IGF-1/PI3K/AKT signalling in aged mice

To assess the exercise-induced modifications in muscle hypertrophy, IGF-1/PI3K/AKT pathway markers were analyzed. The expression levels of IGF-1, PI3K, and AKT in the gastrocnemius muscle after 8 weeks of endurance exercise are shown in Fig 4. The expression level of IGF-1 was significantly higher in Y-Exe mice compared to Y-Con mice (p = 0.023), and it was also significantly higher in A-Exe mice compared to A-Con mice (p = 0.001). Moreover, exercise intervention significantly increased p-AKT/t-AKT expression in both young and aged mice (*p* < 0.049). In addition, the expression level of p-AKT/t-AKT was significantly higher in A-Exe mice than in Y-Exe and A-con mice. In contrast, p-PI3K/t-PI3K levels showed significant differences between groups only in A-Exe (Y-Con, *p* = 0.040; Y-Exe, *p* = 0.042; A-Con, *p* = 0.037).

**Fig 4.**
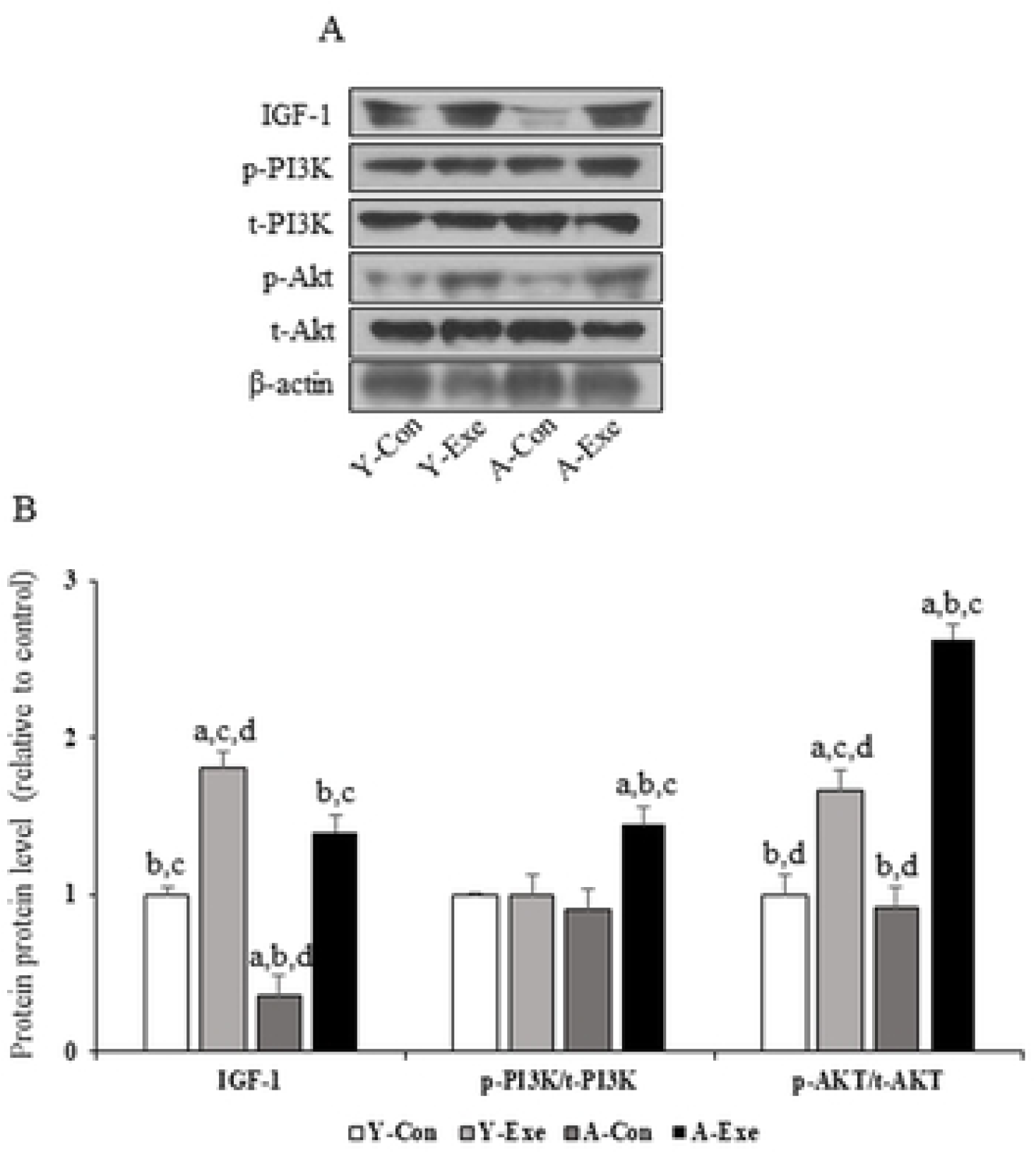
Representative western blots (A), densitometry of IGF-1, p-PI3K, t-PI3K, p-AKT, t-AKT, β-actin (B) in Gastrocnemius tissue from young or old C57BL/6 mice. Y-Con; Young-Control, Y-Exe; Young-Exercise, A-Con; Aged-Control, A-Exe; Aged-Exercise. (a) *p* < 0.05 vs Y-Con, (b) *p* < 0.05 vs Y-Exe, (c) *p* < 0.05 vs A-Con, (d) *p* < 0.05 vs A-Exe, Values are presented a mean±SD; n= 10 per each group.

With respect to exercise-induced muscle cell differentiation in the gastrocnemius muscle, markers of mTOR/p70S6K pathway were assessed (Fig 5). We also showed that 8 weeks of exercise intervention increased the expression of p-mTOR/p-mTOR in both young and aged mice. Further, the level of t-mTOR/p-mTOR was significantly lower in young mice than in aged mice (*p* < 0.018) with A-Exe mice exhibiting the highest level. The expression level of p-p70S6K/t-p70S6K was the highest in A-Exe mice and A-Exe mice showed a significantly higher expression level compared to A-Con mice (*p* < 0.001). Regarding the level of p-p70S6K/t-p70S6K aged mice displayed higher expression levels than young mice, but exercise intervention had no significant effect in either group. Regarding myogenesis, exercise intervention significantly increased the expression of Pax3 (*p* = 0.037) and Myo D (*p* = 0.001) in aged mice and Pax7 (*p* = 0.046) in young mice, while there were significant reductions in Pax3 and MyoD levels in A-Con mice compared to Y-Con mice. Additionally, exercise intervention resulted in a downregulation of Pax3 in young mice and had no significant effects on Pax7 in aged mice and MyoD in young mice (Fig 6).

**Fig 5.**
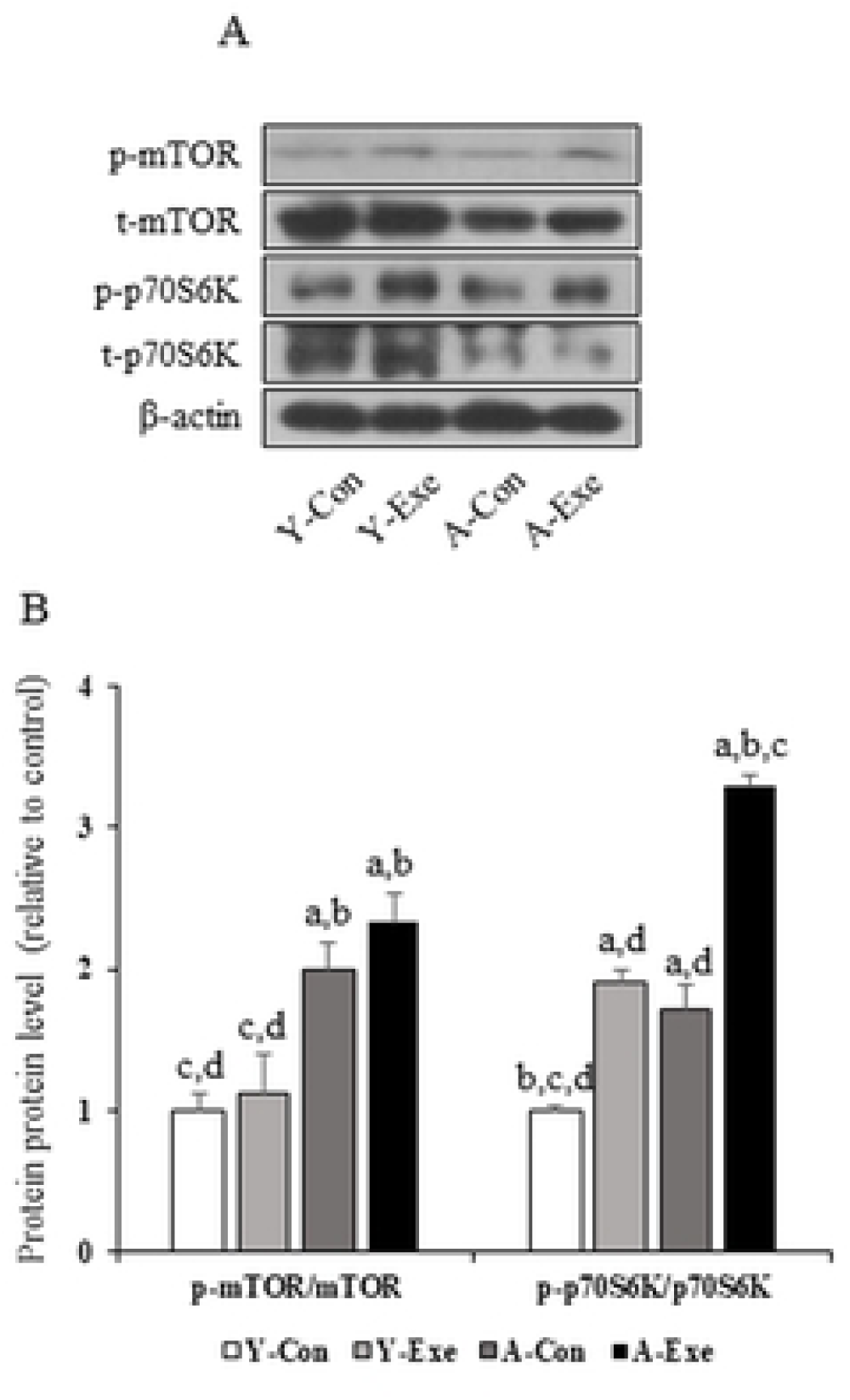
Representative western blots (A), densitometry of p-mTOR, t-mTOR, p-p70S6K, p70S6K and β-actin (B) in Gastrocnemius tissue from young or old C57BL/6 mice. Y-Con; Young-Control, Y-Exe; Young-Exercise, A-Con; Aged-Control, A-Exe; Aged-Exercise. (a) *p* < 0.05 vs Y-Con, (b) *p* < 0.05 vs Y-Exe, (c) *p* < 0.05 vs A-Con, (d) *p* < 0.05 vs A-Exe, Values are presented a mean±SD; n= 10 per each group.

**Fig 6.**
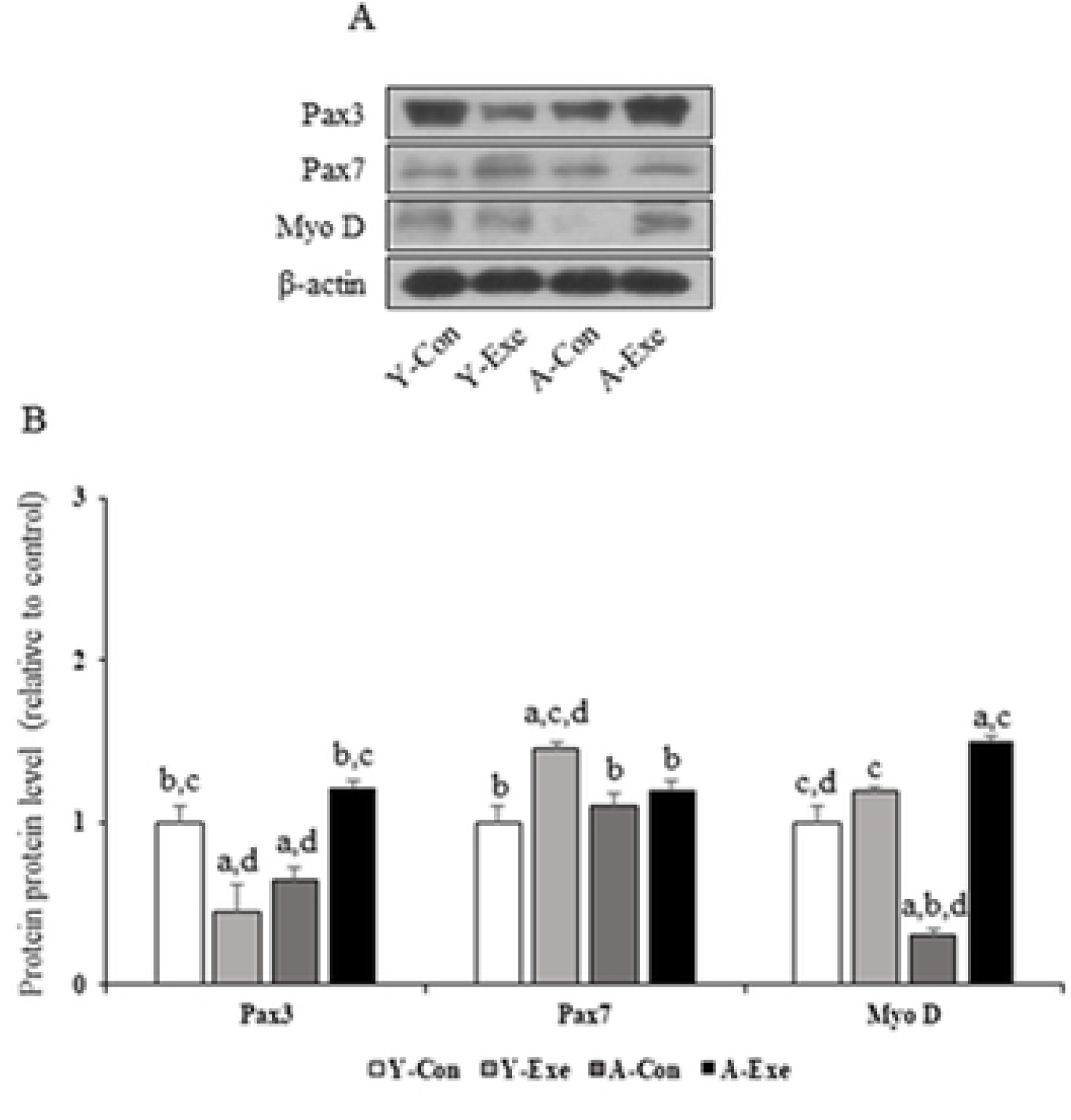
Representative western blots (A), densitometry of Pax3, Pax7, Myo D and β-actin (B) in Gastrocnemius tissue from young or old C57BL/6 mice. Y-Con; Young-Control, Y-Exe; Young-Exercise, A-Con; Aged-Control, A-Exe; Aged-Exercise. (a) *p* < 0.05 vs Y-Con, (b) *p* < 0.05 vs Y-Exe, (c) *p* < 0.05 vs A-Con, (d) *p* < 0.05 vs A-Exe, Values are presented a mean±SD; n= 10 per each group.

## Discussion

The global rise in the elderly population poses significant challenges, spanning healthcare needs to social and economic implications. With the increasing number of older individuals, there is a corresponding surge in musculoskeletal disorders such as osteoporosis, arthritis, and sarcopenia, which can markedly affect mobility and quality of life. Regular exercise has been shown to mitigate the effects of aging on muscles by promoting muscle growth, enhancing strength, and improving endurance, thus enhancing overall physical function and promote independence among older adults. Consequently, this study was conducted to investigate the effects of 8 weeks of exercise training on the progressive reduction in muscle mass and function in aged mice.

### Effect of 8 weeks of exercise intervention on muscle strength and muscle fiber size

Muscle atrophy, a common phenomenon in the elderly, has significant implications for independent living and overall quality of life ^22^. Longitudinal studies, such as that conducted by Goodpaster et al. ^23^, have observed declines in muscle mass and strength among over 1800 elderly individuals aged over 70 years during a 3-year follow-up period. Similarly, research by Miller et al. ^24^ and Aversa et al. ^25^ has corroborated age-related losses in muscle mass and diminished muscle performance. Animal studies, such as those conducted by Wang et al. ^26^ and Muller et al. ^27^ have provided insights into the mechanisms underlying muscle atrophy and oxidative stress associated with aging. These findings align with the present study, where observations revealed decreased muscle fiber size and strength in aged mice compared to their younger counterparts.

In response to these age-related changes, interventions such as endurance exercise have emerged as potential strategies to mitigate muscle decline. The current study demonstrated that 8 weeks of endurance exercise led to increased muscle strength and fiber size in the gastrocnemius muscle of both young and aged mice. These findings are consistent with prior research suggesting that exercise training can prevent muscle atrophy, preserve muscle strength, and facilitate recovery from age-related muscle fiber damage ^28, 29^. Furthermore, studies on elderly populations have shown promising results regarding the benefits of strength training exercises. Aas et al. ^30^ found that 10 weeks of lower-body strength exercise led to significant increases in muscle size among 85-year-old individuals. Similarly, Lee et al. ^31^ reported improvements in muscle strength and mass among community seniors following band exercises. Animal studies, such as the one conducted by Zhao et al. ^32^, have further supported the efficacy of exercise interventions in preserving muscle health. Their research demonstrated that moderate-intensity exercise intervention attenuated decreases in muscle mass, muscle fiber cross-sectional area, and myonuclei numbers in mice aged between 24 to 56 weeks. White et al. ^29^ also demonstrated that resistance wheel exercise had a significant impact on mitochondrial and autophagosomal pathway, ultimately mitigating age-related muscle atrophy.

### Effect of 8 weeks of exercise intervention on motor performance

As aging progresses, the functional capacity of muscle gradually diminishes, leading to decreased exercise performance. This decline in physical function has been associated with an increased risk of falls, fractures, metabolic diseases, and mortality among older adults ^33^. Wages et al. ^34^ conducted a comprehensive evaluation of physical function in elderly individuals aged 70 years and older, assessing grip strength, thigh extensor strength, 6-minute walking speed, and stair climbing ability. Their findings revealed significant correlations between grip strength, femoral extensor strength, walking speed, and chair climbing time, highlighting the importance of muscle strength in maintaining mobility and functional independence. Moreover, Liu et al. ^35^ demonstrated that older adults with higher arm strength exhibited better performance on dexterity control and hand function tests compared to those with lower grip strength. This decline in motor performance among the elderly may stem from impairments in the nervous system, leading to reduced motor coordination, increased movement variability, and balance and gait disorders ^36–38^.

Consistent with these findings, our study observed lower motor coordination in aged mice compared to younger counterparts in the Rotarod test. However, exercise training interventions improved motor coordination in both young and aged mice, aligning with previous research. For instance, Shahtahmassebi et al. ^39^ implemented a 12-week exercise program in elderly individuals aged 60 years and older, resulting in increased muscle fiber size in the rectus abdominis and lumbar multifidus muscles. Similarly, Michaelson et al. ^40^ reported that exercise interventions significantly enhanced muscle strength and cross-sectional area, thereby reducing the incidence of falls in elderly individuals aged 91 years. Furthermore, exercise training has been shown to improve muscle strength and exercise performance across various functional assessments, including the Performance Oriented Mobility Assessment, Physical Performance Score, Groningen Activity Restriction Scale ^41^, timed up and go test, and 10-Meter Walk Test ^42^. Collectively, these findings underscore the critical role of exercise in enhancing the quality of life and preserving functional independence among older adults.

### Effect of 8 weeks of exercise intervention on the IGF1/PI3K/AKT pathway

Considering the IGF1 signaling pathway, the current findings showed that the expression levels of IGF-1, p-AKT, p-mTOR, t-mTOR, p70S6K, and p-p70S6K were decreased in aged mice, indicating muscle atrophy. This observation aligns with previous research indicating a decline in the expression levels of AKT, mTOR, and p70S6K with aging in skeletal muscle ^43^. Moreover, aging has been associated with a downregulation of IGF-1 expression both systemically and locally, further exacerbating muscle atrophy ^44^. The diminished expression of IGF-1 is often accompanied by an increase in insulin-like growth factor-binding proteins, which bind to IGF-1, reducing its bioavailability and activity in peripheral tissues. Conversely, elevated IGF-1 expression has been shown to mitigate muscle atrophy associated with aging in murine models ^45^. Additionally, IGF-1 isoforms such as IGF-1Ea and IGF-1Eb also have been demonstrated protective effects against age-related muscle loss and force decline. They can stimulate muscle growth, trigger the autophagy/lysosome mechanisms and boost PGC1-α level ^19^.

Remarkably, our study demonstrated that exercise training effectively activated the IGF1/phosphoinositide 3-kinase (PI3K)/AKT pathway. This finding aligns with previous investigations. For instance, Chen et al. ^46^ implemented various exercise regimens, including resistance, aerobic, and complex training programs, in elderly individuals aged 65 to 75 years, revealing an increase in serum IGF-1 expression in the exercise groups. Similarly, Yoon et al. ^47^ observed improvements in sleep quality following 12 weeks of resistance and endurance exercises in elderly individuals aged 64 years or older. Zeng et al. ^48^ reported a significant upregulation of mRNA expression levels of IGF-1, as well as activation of mTOR, p70S6K, and eukaryotic translation initiation factor 4E-binding protein 1 (4EBP1), in response to exercise interventions. Furthermore, Yin et al. ^49^ demonstrated that resistance and aerobic exercises induced enhancements in the levels and activities of IGF-1, IGF-1 receptor, mTOR, PI3K, and AKT, resulting in augmented muscle hypertrophy in male Sprague Dawley rats.

The study faced several limitations. Firstly, the sample size was relatively small, potentially limiting the generalizability of the findings to larger populations. Secondly, there was no attempt to quantify age-related loss of motor neurons and muscle fibers. Lastly, the visual images displayed tears and cracks, possibly due to dehydration during the H&E staining process, highlighting the need for improved visual evidence in future studies.

## Conclusion

Overall, this study investigated the effect of 8 weeks of exercise intervention on age-related musculoskeletal atrophy and decreased motor performance in age mice and revealed that exercise intervention 1) increases grip strength in aging mice; 2) increases the muscle size in aged mice, though the differences are not statistically significant; 3) improves motor performance in aging mice; and 4) increases the expression of muscle fiber growth and regeneration markers in the gastrocnemius muscles of aged mice. Overall, regular exercise improves various signaling pathways related to muscle strength and muscle fiber growth, as well as prevents various degenerative diseases and improves the quality of life in the elderly.

## Acknowledgments

This work was supported by the Ministry of Education of the Republic of Korea and the National Research Foundation of Korea (NRF-2020S1A5B5A16083059)

## Human and Animal Rights

All experiments and procedures were approved by the Institutional Animal Care and Use Committee of Sungkyunkwan University

